# Trans-epithelial migration is essential for neutrophil activation during RSV infection

**DOI:** 10.1101/2021.10.04.463016

**Authors:** Elisabeth Robinson, Jenny Amanda Herbert, Machaela Palor, Luo Ren, Isobel Larken, Alisha Patel, Dale Moulding, Mario Cortina-Borja, Rosalind Louise Smyth, Claire Mary Smith

## Abstract

The recruitment of neutrophils to the infected airway occurs early following respiratory syncytial virus (RSV) infection and high numbers of activated neutrophils in airway and blood is associated with the development of severe disease. Here, we investigated whether trans-epithelial migration across primary human airway epithelial cells (AECs) is sufficient and necessary for neutrophil activation during RSV infection. Using flow cytometry, we identified three populations of neutrophils in our *in vitro* model; those in suspension in basolateral and apical compartments and those that migrated and adhered to AECs. After 1h incubation, the number of adherent neutrophils was significantly greater following RSV infection compared to mock infected. We found that, when migration occurred, neutrophil expression of CD11b, CD62L, CD64, NE and MPO increased in all compartments. However, this did not occur when neutrophils were prevented from migrating. This suggests that the heightened neutrophil activation we detected in the basolateral compartment may be due to reverse migrating neutrophils, as has been suggested by clinical observations. Using live-cell fluorescent microscopy, we then profiled the early temporal and spatial movement and adherence of human neutrophils during migration. Our findings suggest three main phases of early neutrophil recruitment and behaviour in the airways during RSV infection, with neutrophil recruitment, activation and adherence to RSV infected AECs, with clustering, occurring within the first 20 minutes. This work and the model we developed could provide new insight into how neutrophil activation and a dysregulated neutrophil response to RSV mediates disease severity.

**Graphical Abstract:** Building on previous work of neutrophil function we propose 3 main phases of early neutrophil recruitment and behaviour in the airways during RSV infection. Phase 1. Initial chemotaxis and adherence: Here unstimulated circulating neutrophils expressing baseline levels of CD11b migrate across infected AECs in response to chemotactic signals in the apical supernatant. Some neutrophils remain adherent to the infected AECs. Phase 2: Activation and reverse migration: once on the apical side of the epithelium, neutrophils increase expression of CD11b and other activation associated markers, and some ‘activated’ neutrophils undergo reverse migration. Neutrophils with greater expression of CD11b are detected on the basolateral side Phase 3: Amplified chemotaxis and clustering: after 20 minutes, adherent neutrophils begin to rapidly cluster on RSV infected primary airway epithelial cells cultures, mediated by signalling from a dying neutrophil. Drawing created using BioRender.com.

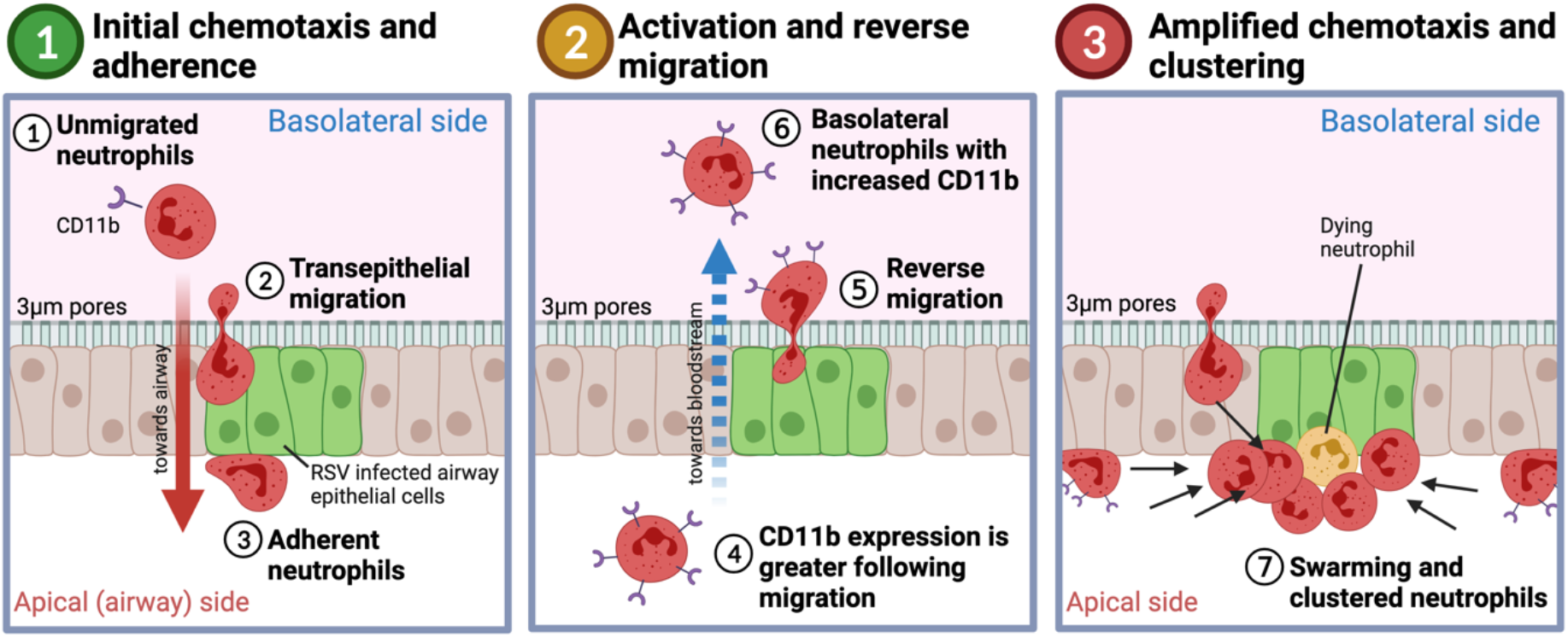

## Introduction

Respiratory Syncytial Virus (RSV) is a seasonal respiratory virus, reported to infect almost all children before the age of 2(1). Following infection, most children develop an illness which is confined to upper respiratory tract symptoms, however 1-3% of infected infants will develop a severe illness requiring hospitalisation(2). There is no licensed vaccine for RSV and treatment is currently limited to supportive care. In resource-limited countries, RSV is a major cause of infant mortality (3–5).

Although risk factors have been identified, it is still not clear why RSV infected infants, with no apparent risk factors, require hospitalisation and respiratory support (6). One suggestion is that neutrophils, which form around 80% of all cells recovered from the airways of infants with severe RSV bronchiolitis by bronchoalveolar lavage(7), and their associated cytokines contribute to disease severity(8–10). Neutrophil mediated factors such as CXCL8 (IL-8), CXCL10 (IP-10) and neutrophil elastase are also present in substantial amounts in airway secretions of children with severe RSV bronchiolitis. (11–13).

Neutrophils are the predominant immune cell type in the systemic circulation and are the first cell recruited from the bloodstream through the airway epithelium during infection(14). Here they are thought to employ largely non-specific mechanisms of pathogen destruction including phagocytosis, neutrophil extracellular trap formation and release of toxic granule products such as neutrophil elastase and myeloperoxidase(15,16). Neutrophil trans-epithelial migration is facilitated by neutrophil receptors including integrins (i.e. CD11b) and selectins (such as CD62L) that bind to molecules on airway epithelial cells such as ICAM-1(17–19). Neutrophils also interact with other immune complexes using Fc receptors such as CD64; upregulation of CD64 has been evaluated as a biomarker for neonatal sepsis in infants(20).

Our previous work has shown that neutrophils migrate across primary differentiated RSV infected AECs resulting in increased epithelial damage and a reduced viral load (21,22). We observed that neutrophils adhere to RSV infected AECs in a clustering pattern, which was not seen in uninfected AECs. This pattern of neutrophil chemotaxis and migration has been reported using intravital microscopy, referred to as ‘neutrophil swarming’ due to a resemblance with the swarming behaviour of insects (23,24). We also showed that migrated neutrophils have greater cell surface expression of CD11B and MPO compared to neutrophils that had not migrated^21^. What remains unclear is whether migrated neutrophils with (i.e. higher CD11b) are selected for trans-epithelial migration or whether they become activated due to migration *per se.* Although RSV is not known establish infection outside the respiratory tract or cause a viraemia, clinical studies have shown that a high proportion of neutrophils from the systemic circulation of infants with RSV bronchiolitis contain RSV mRNA(11). This raises the possibility that activated neutrophils, recruited to the airway during RSV infection, may be able to migrate in the reverse direction, back to the systemic circulation.

This study investigates the behaviour and function of neutrophils as they move across the airway epithelial cell layer during RSV infection. We studied this migration using a human model of primary airway epithelial cells grown at air-liquid interface for 7 days. This model facilitated the higher-resolution z-stack microscopy needed for 4D (XYZT) tracking of neutrophil migration for the first time. (**Figure 1**). We used this model evaluate the kinetics of neutrophil trans-epithelial migration including ‘cluster formation’, and the temporal and spatial association of neutrophil activation during RSV infection.

**Figure 1.**
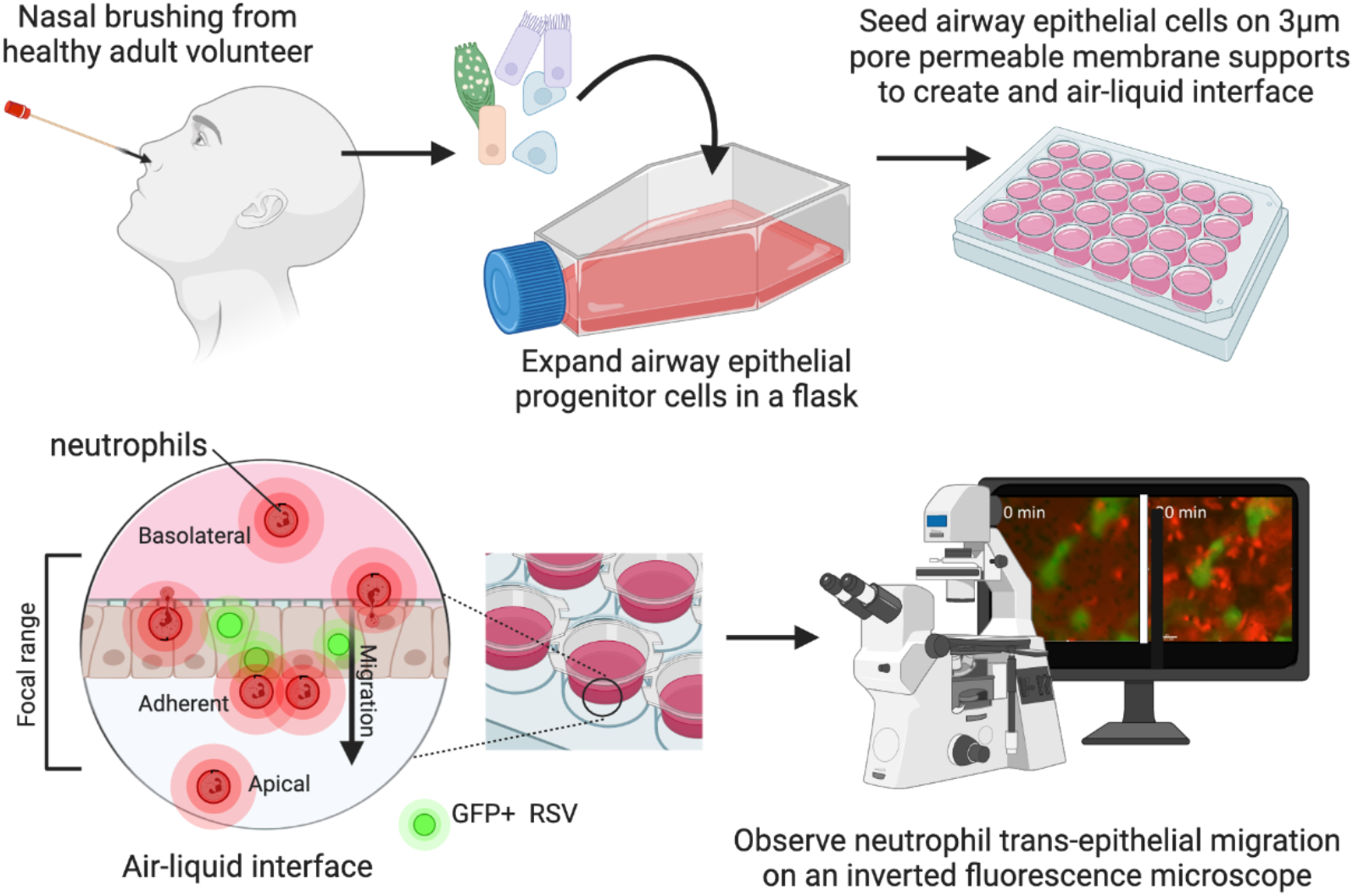
Schematic of method and model used to study neutrophil trans-epithelial migration in response to RSV infection. Human primary airway epithelial cells (AECs) were cultured on the underside of a 3μm pore size, transparent PET culture membrane insert. AECs were matured at an air-liquid interface (ALI) for 7 days before infection with GFP tagged RSV. At this time, neutrophils, stained with a viability stain (calcein-red orange), were added to the basolateral side of AEC cultures and a 50μm Z-stack image of the focal area indicated was captured for up to 1hr. Drawing created using Biorender.com

## Results

### Neutrophil trans-epithelial migration increases damage to airway epithelial cells associated with a release of neutrophil proteases

Firstly, as we have done before in differentiated cultures (22), we characterised whether neutrophil migration increases epithelial damage during RSV infection in our novel AEC model. As controls, we compared epithelial damage and neutrophil protease release across either 1) mock infected AECs, 2) mock infected AECs exposed to potent neutrophil chemoattractant (fMLP), or 3) mock infected AECs exposed to RSV infected AEC supernatant (referred to as RSV Sup). These controls allowed us to differentiate whether our observations were due to the process of migration *per se* or to inflammatory mediators present in the RSV infected airway supernatant.

We found that 1h after neutrophil trans-epithelial migration across epithelium infected with RSV for 24h, we detected larger gaps with mean ± standard error of the mean (SEM) of 70.8 ± 4.6% area in the RSV infected epithelial layer compared to the mock-infected (61.6 ± 6.0% area) (*p*<0.0001) (representative images shown in **Figure 2A)**. We also detected a loss of RSV infected cells, as is observed in **Video 1 (video stills shown in video stills A).** Here as neutrophil transepithelial migration progresses, an AEC expressing GFP RSV indicated by a yellow circle, disappears from the image. A video of neutrophils migrating across mock infected cells is shown in **Video 2.**

**Figure 2.**
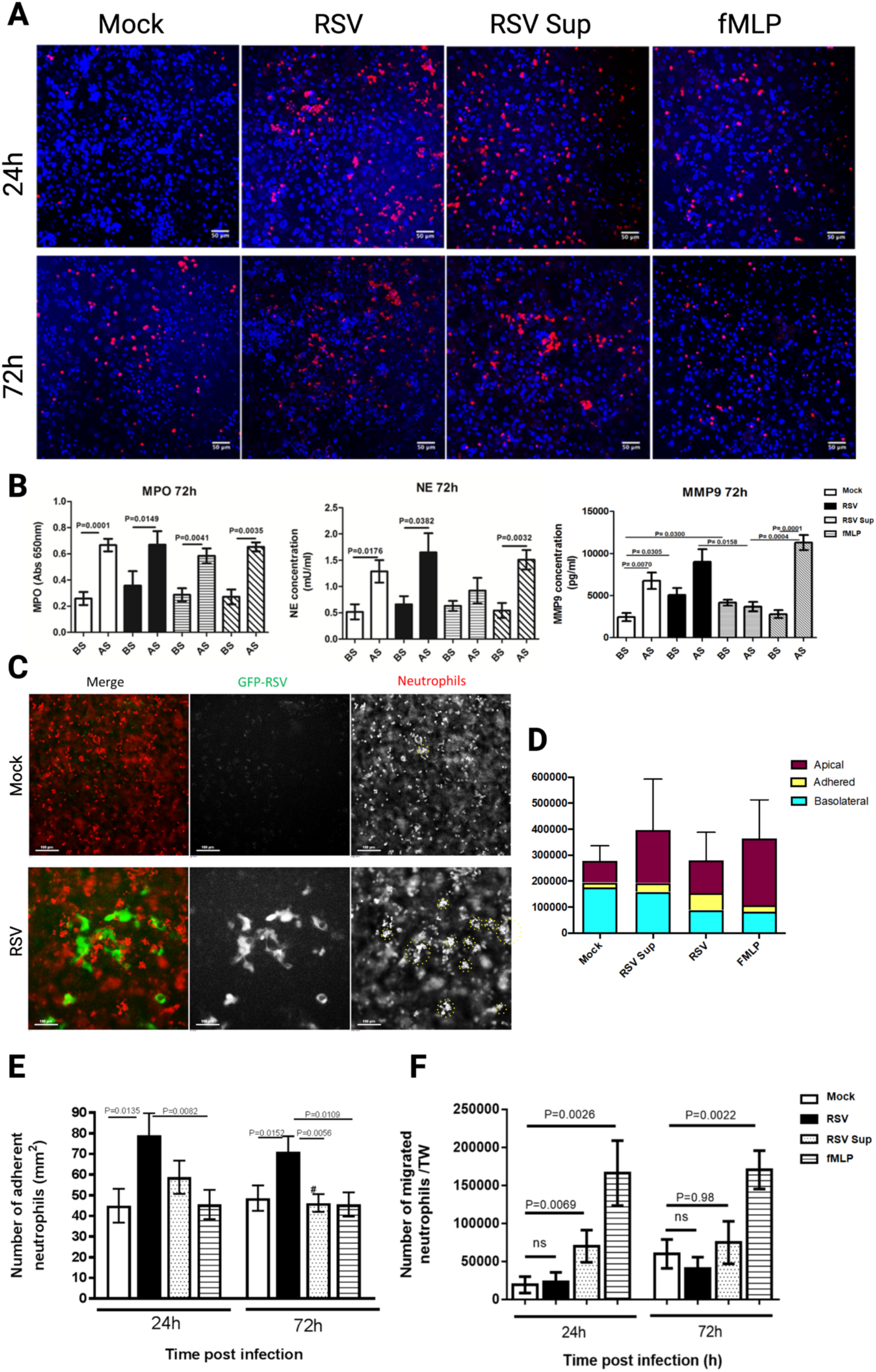
(Overleaf) RSV increases the numbers of neutrophils that adhere to RSV infected epithelial cultures, forming clusters on the AEC surface. **(A)** – Representative image showing neutrophil (red cells) adherence to epithelial cells (DAPI staining – blue) following migration for 1h at 24h or 72h post RSV or mock infection. fMLP 1ng/ml placed apical to uninfected AECs was used a positive control for neutrophil chemotaxis. Scale bar indicates 50μm (**B**) – Soluble granular factors myeloperoxidase (MPO), neutrophil elastase (NE) and matrix-metalloproteinase-9 (MMP-9) in the apical supernatant (designated AS in figure) and basolateral supernatant (BS) were quantified using ELISA. fMLP and mock infected AECs with supernatant collected from RSV infected AECs placed apically (RSV Sup). (**C**) Representative 2 channel maximum intensity projection of a Z-stack image (50μm range) of 72-hour mock (top) and RSV (bottom) showing neutrophil adherence in clusters to mock and RSV infected AECs. Green = gfpRSV infected AECs, red=neutrophils. Scale bar indicates 100μm (**D**) – Graph showing representative counts of neutrophils isolated either basolateral to, apical to or adherent to AECs. Absolute counts performed using a flow cytometer selecting for neutrophils as positive for CD11b-APC antibody staining. (**E**)– Numbers of neutrophils adherent to AECs after 1hr transepithelial migration across 24 or 72 hr RSV infected AECs. Adherent neutrophils and epithelial cells were counted using ImageJ counting tool, the average number of neutrophils from all images is shown. (**F**)-Numbers of neutrophils migrating and dissociating apically from AECs after 1hr transepithelial migration across 24 or 72 hr RSV infected AECs. Neutrophil concentrations were quantified in the apical surface media using a plate reader and read against a standard curve.

**Video Stills A.**
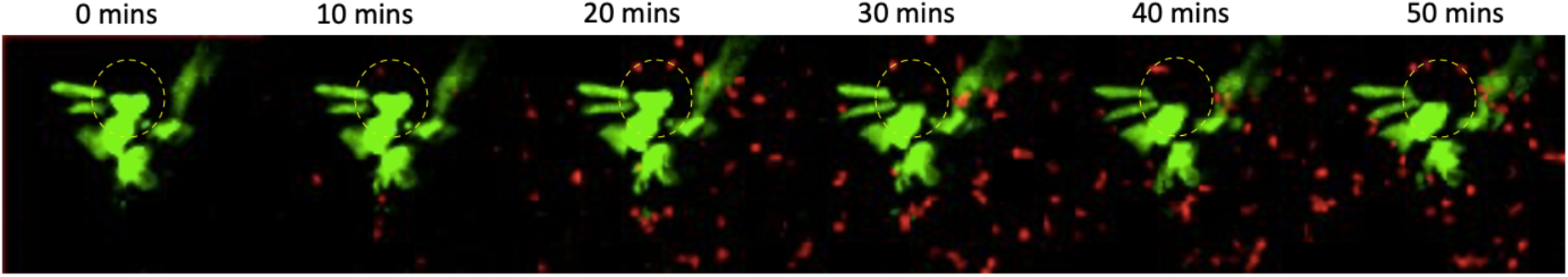
Elimination of RSV infected AEC during neutrophil migration through RSV infected AECs. Representative stills of a time lapse stack detailing neutrophil migration through RSV infected AECs. Green indicates RSV infected cells expressing GFP and red indicates neutrophils stained with calcein red-orange. Yellow dashed circle shows position of RSV-infected epithelial cell.

Neutrophils release several toxic products, including myeloperoxidase (MPO) and neutrophil elastase (NE). We measured the concentration of these products in the compartments of our model after neutrophil migration. We found that the concentration of NE, in airway surface media of AEC cultures, was >3 fold greater than the basolateral media following neutrophil migration across RSV infected epithelium for 4h at 72h post infection (**Figure 2B**) with a mean ± SEM of 2.0 ± 0.6mU/ml, compared to 0.6 ± 0.1mU/ml in the mock-infected cultures (*p*=0.039) (**Figure 2B**). We did not find a significant difference in MPO in airway surface media from RSV infected cultures following neutrophil migration for 4h compared to the mock-infected cultures.

### RSV infection results in increased neutrophil adherence to AECs

As we previously found that neutrophil adherence was associated with epithelial damage to RSV infected ciliated cultures (22), here we also we counted the number of neutrophils that migrate across and adhere to AECs using fluorescence microscopy (**Figure 2C**). Overall, transepithelial migration did not affect neutrophil viability (**Figure 2D**). The total counts of viable neutrophils per well, i.e. combined basolateral, apically and adherent neutrophils, is shown (**Figure 2D**). The numbers of viable neutrophils adherent to AEC cultures infected with RSV for 24h (791.1 ± 106.8 cells/cm^2^) or 72h (711.1 ± 74.3 cells/cm^2^) were significantly (*p* = 0.014) greater than the respective mock infected AEC cultures (449.8 ± 81.82 cells/cm^2^) or (486.6 ± 61.5 cells/cm^2^), respectively (**Figure 2E**).

However, at 72h post infection, significantly (p = 0.006) fewer neutrophils remained adherent to mock infected AECs exposed to RSV infected AEC supernatant (462.7 ± 43.3 cells/cm^2^), in comparison to the RSV infected AECs (711.1 ± 74.3 cells/cm^2^) (Figure 2E). This implies that neutrophil adherence is facilitated by RSV infected epithelial cells, rather than soluble, secreted factors released by RSV infected AECs alone. This was supported by our finding that RSV infection did not lead to an increase in the number of apical neutrophils (those that migrate and detach from the epithelial cells), with an average (mean ± SEM) of 22,876 ± 12,713 neutrophils/well, compared to the mock infected AEC cultures (19,184 ± 10,806) at 24h post infection (Figure 2F). Whereas significantly (p = 0.007) more apical neutrophils were recovered from mock infected AECs exposed to RSV infected AEC supernatant (RSV Sup), with an average 70,016 ± 21,115 neutrophils/well compared to the mock with 19,184 ± 10,806 and 22,876 ± 12,713 in comparison to RSV (p = 0.006) (Figure 2F). These data were similar at 72 hours post-infection (p = 0.98) the number of neutrophils recovered from the RSV infected AEC cultures (40,415 ± 15,143 neutrophils/well) compared to the mock (59,900 ± 18,885) (Figure 2F).

### Neutrophils upregulate expression of key surface markers following migration across AECs

So far, this study has shown that neutrophils are capable of trans-epithelial migration across airway epithelial cells and either ‘remain’ on the *basolateral* side, become *adherent* to AECs, or migrate and dissociate into the *apical* space (see **Figure 1**). To determine whether the ability of a neutrophil to migrate and adhere to epithelial cells is associated with the expression of specific cellular markers associated with neutrophil activation and migration (i.e., CD11b, CD64, CD62L, NE, MPO), we analysed neutrophils recovered from basolateral, adherent, and apical compartments using flow cytometry. We chose the 72-hour timepoint following AEC infection, and 1h timepoint after neutrophil migration for these experiments as these were the conditions that resulted in the greatest number of adherent neutrophils (**Supplementary Figure 1E**) to allow effective comparisons.

Firstly, as a control, we determined whether naïve neutrophils altered their expression of CD11b, CD64, CD62L, NE, MPO following exposure to apical supernatants recovered from mock or RSV infected AECs, but in the absence of AECs (**Figure 3** see “NoAEC”). Here, we found a significantly greater expression of CD11b on neutrophils incubated with RSV infected AEC supernatant (4,078.3 ± 109.8) compared to neutrophils incubated with media alone (1,583.7 ± 34.3) (*p*<0.001) or mock infected AEC supernatant (2,751 ± 37.5) (*p*=0.001) (**Figure 3A**). However, we found no significant difference in CD62L, CD64, NE or MPO expression on neutrophils incubated with supernatants from mock and RSV infected AECs (**Figure 3BCD&E**).

**Figure 3.**
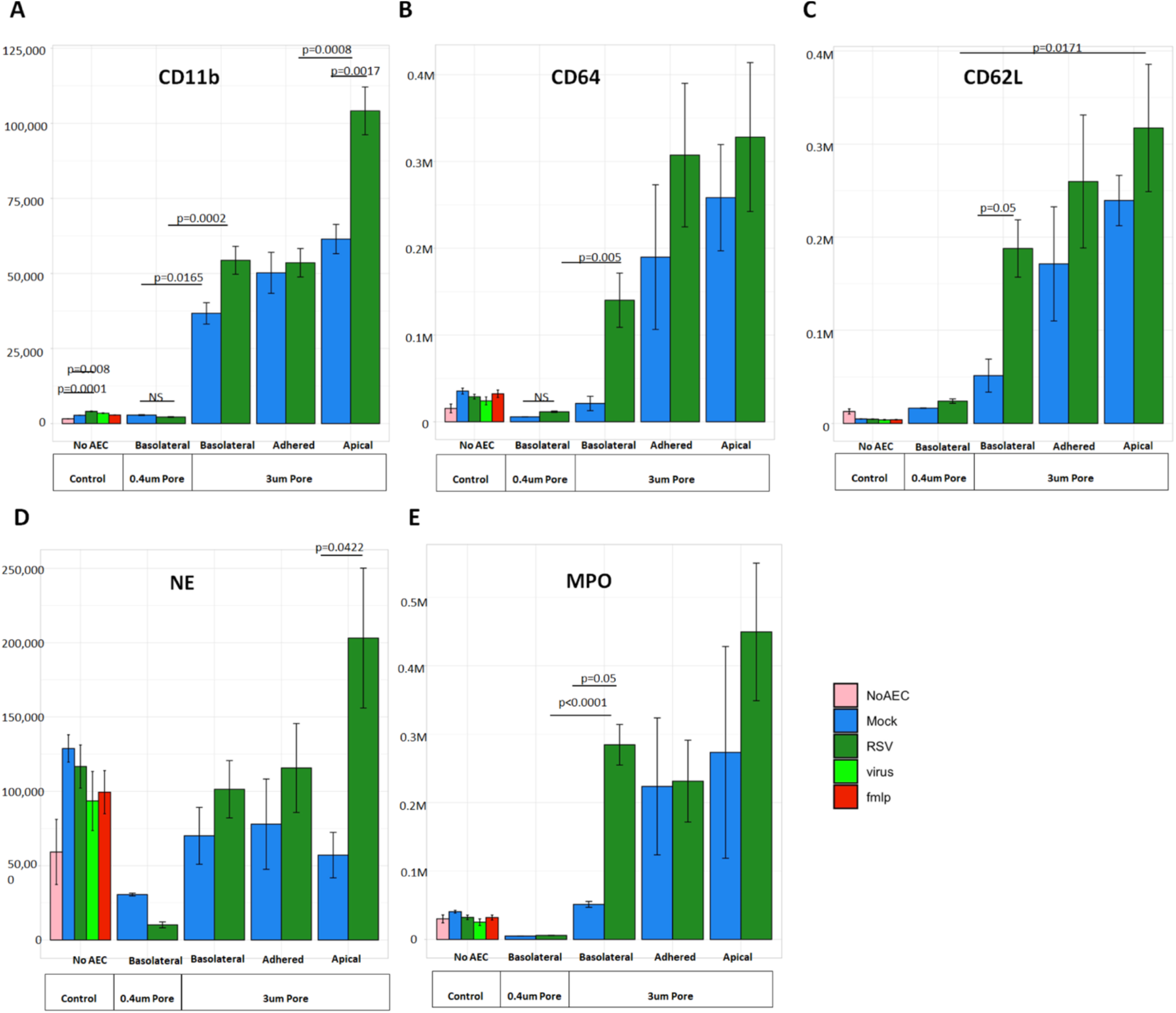
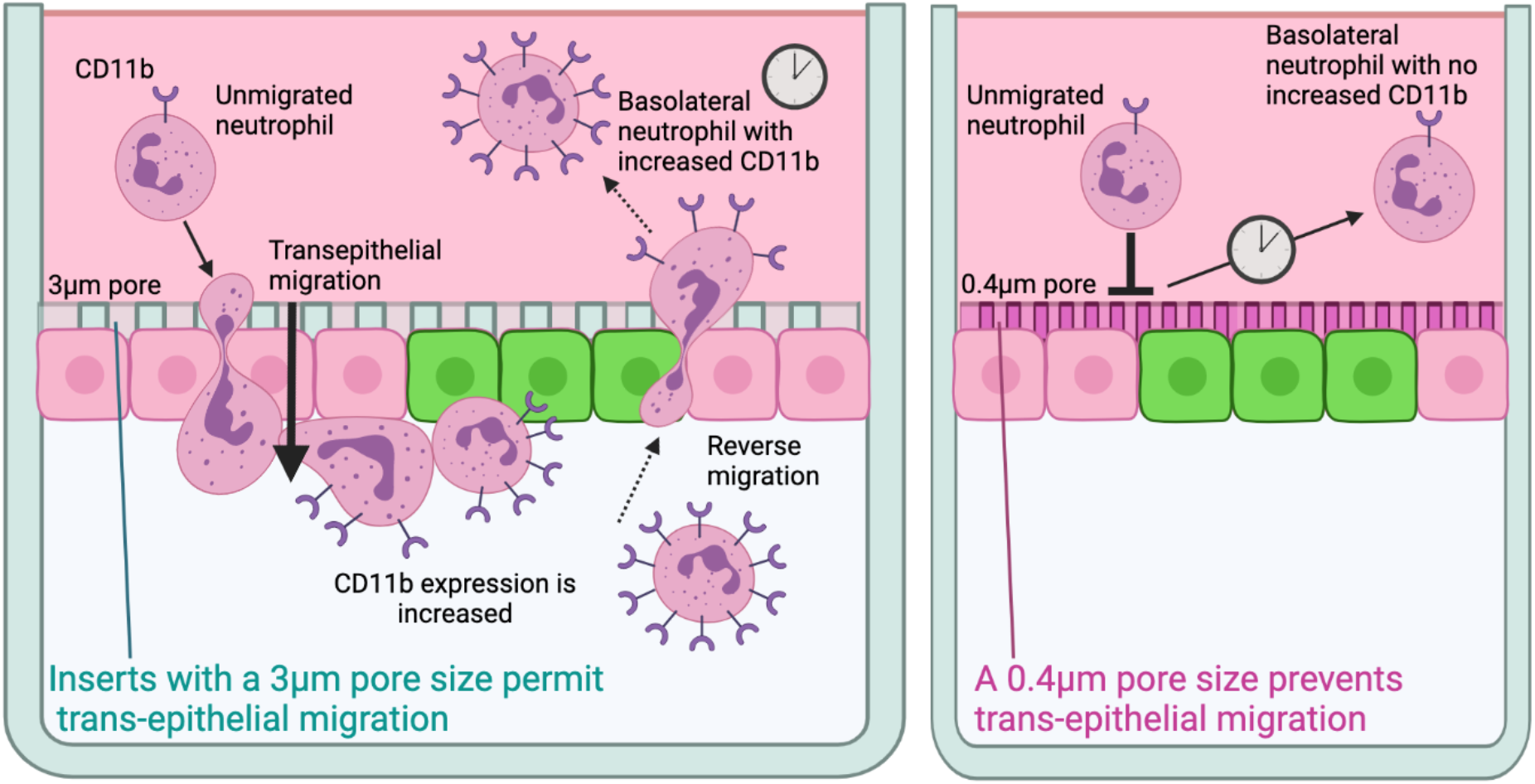
Neutrophils undergoing trans-epithelial migration alter their expression of surface markers depending on their location. **(A-E)** Mean fluorescence intensity of cell surface expressed CD11b (A), CD64 (B), CD62L (C), NE (D), MPO (E) on neutrophils incubated for 1h with media alone (No AEC) or following migration across mock or RSV infected AEC cultures infected for 72 hours on membrane inserts with a 3μm or 0.4μm pore size, the latter prevents cellular migration (see **Supplementary Figure S2)**. Graphs indicate neutrophils recovered from the basolateral, adherent, and apical (migrated) compartments within the assay. Neutrophils incubated with fMLP and RSV MOI1 (virus) in media without AEC were also used as controls. Bars indicate standard error of the mean. A linear model of mixed effects was used to compare interactions between infection and location groups and control for intra-donor variability. Individual comparisons performed with Two-Way ANOVA with pairing and Tukey’s post hock test. * = in comparison to respective basolateral group # = in comparison to respective adherent group. (**F**)- Graphical illustration of possible interpretation of findings of CD11b expression **Left panel** shows primary airway epithelial cells cultured at ALI on membrane inserts with 3μm pore size that <u>permits</u> neutrophils to migrate through. 1) unmigrated neutrophils expressing baseline levels of CD11b, 2) neutrophils migrate across infected AECs and some remain adherent to the infected AECs, 3) neutrophils are shown to increase expression of CD11b and other activation associated markers, 4) some ‘activated’ neutrophils undergo reverse migration as 5) neutrophils with the increased expression of CD11b are detected on the basolateral side of the insert. **Right panel** shows primary airway epithelial cells cultured at ALI on membrane inserts with 0.4μm pore size that <u>prevents</u> neutrophil contact with AECs and migration. Here, after 1h incubation with RSV infected AECs, neutrophils on the basolateral side 1) and 2) were shown to express the same level of CD11b markers, indicating that neutrophil contact with AECs and starting to move through the AECs is key process for increasing expression of these activation markers. Drawing created using BioRender.com.

Introducing neutrophils to our AEC model led to a significant increase in expression of CD11b and CD64 (*p* = 0.0002, *p* = 0.005) in neutrophils recovered from basolateral, adherent, apical compartments of RSV infected AEC models compared to neutrophils exposed to media alone (non-AEC control) (**Figure 3ABC**) (comparison not directly shown on graph). Interestingly, we detected a significant (*p* = 0.0076) 24 and 30-fold increase (*p* <0.0001) in CD11b expression on neutrophils recovered from the *basolateral* compartments of both mock (36,708 ± 3,563) and RSV infected AECs (54,389 ± 3,863) compared to neutrophils exposed to media alone (non-AEC control) (1,583.7 ± 34.3) (**Figure 3ABC**). To determine whether this was due to factors secreted into the basolateral compartment by the AECs, and therefore independent of migration, AECs were grown on membranes with a 0.4μm pore size. This smaller pore size does not permit neutrophils to move across the membrane and have contact with AECs (**Supplementary Figure S2**) but will allow for passive diffusion of secreted factors. Using this system, we found that the expression of CD11b, CD64, CD62L and MPO on basolateral neutrophils exposed to AECs grown on inserts with a 0.4μm pore size did not increase and, in fact, was no different to the values obtained in the absence of AECs. Compared to neutrophils recovered from the basolateral side of the RSV infected AEC cultures grown on 0.4μm inserts, demonstrated CD11b, CD64, CD62L, NE and MPO levels that were at least 30x lower compared to those incubated on AECs grown on inserts with 3μm pores (**Figure 3**). This reduction in neutrophil activation was significant (*p* < 0.05) across all groups tested (mock and RSV infected) suggesting the increased expression of these markers in the basolateral and adhered neutrophils is a result of direct contact with the AECs rather than infection (**Figure 3F**).

Following trans-epithelial migration, we found that RSV infection was associated with a further 1.5-fold increase (*p* = 0.021) in CD11b expression on apical (104,145 ± 6,631) neutrophils compared to neutrophils recovered from the respective compartments of mock infected cultures (61,466 ± 4,876) (see **Figure 3A**). There was no significant difference in CD64, CD62L, NE, MPO expression on neutrophils recovered from RSV compared to mock infected AECs (**Figure 3BCD&E**). The highest expression levels for CD11b were recorded on apical neutrophils recovered from RSV infected AECs (104,145 ± 6,631), which was more than two-fold higher (*p* < 0.05) than both basolateral neutrophils (54,389 ± 3,863) and adherent neutrophils (53,561 ± 3,932) in the same model and >10,000-fold higher than neutrophils in media alone (no-AEC) (1,583.7 ± 34.3) (**Figure 3A**), suggesting that both RSV infection and migration of neutrophils was associated with this high expression of CD11B.

### Temporal analysis of neutrophil trans-epithelial migration reveals that migrated neutrophils move slower but farther during RSV infection

We used time-lapse fluorescence microscopy imaging and performed vector analysis (or XYZ coordinates) of fluorescently labelled neutrophils to determine differences in the speed, track duration, distance travelled and directionality of neutrophils (n=200 per condition) during trans-epithelial migration through mock and RSV infected AECs.

#### SPEED

We found that, over the course of the first hour, neutrophils’ average (mean ±SEM) speed across RSV infected AECs was significantly (p<0.001) slower (2.28μm/sec ± 0.008) than the average speed of neutrophils moving through the mock infected AECs (4.18μm/sec ± 0.14) (**Figure 4A)**. The exception to this was the first 15 minutes of exposure, where we found no significant difference in speed of neutrophils migrating across RSV and mock infected AECs (**Figure 4A)**.

**Figure 4.**
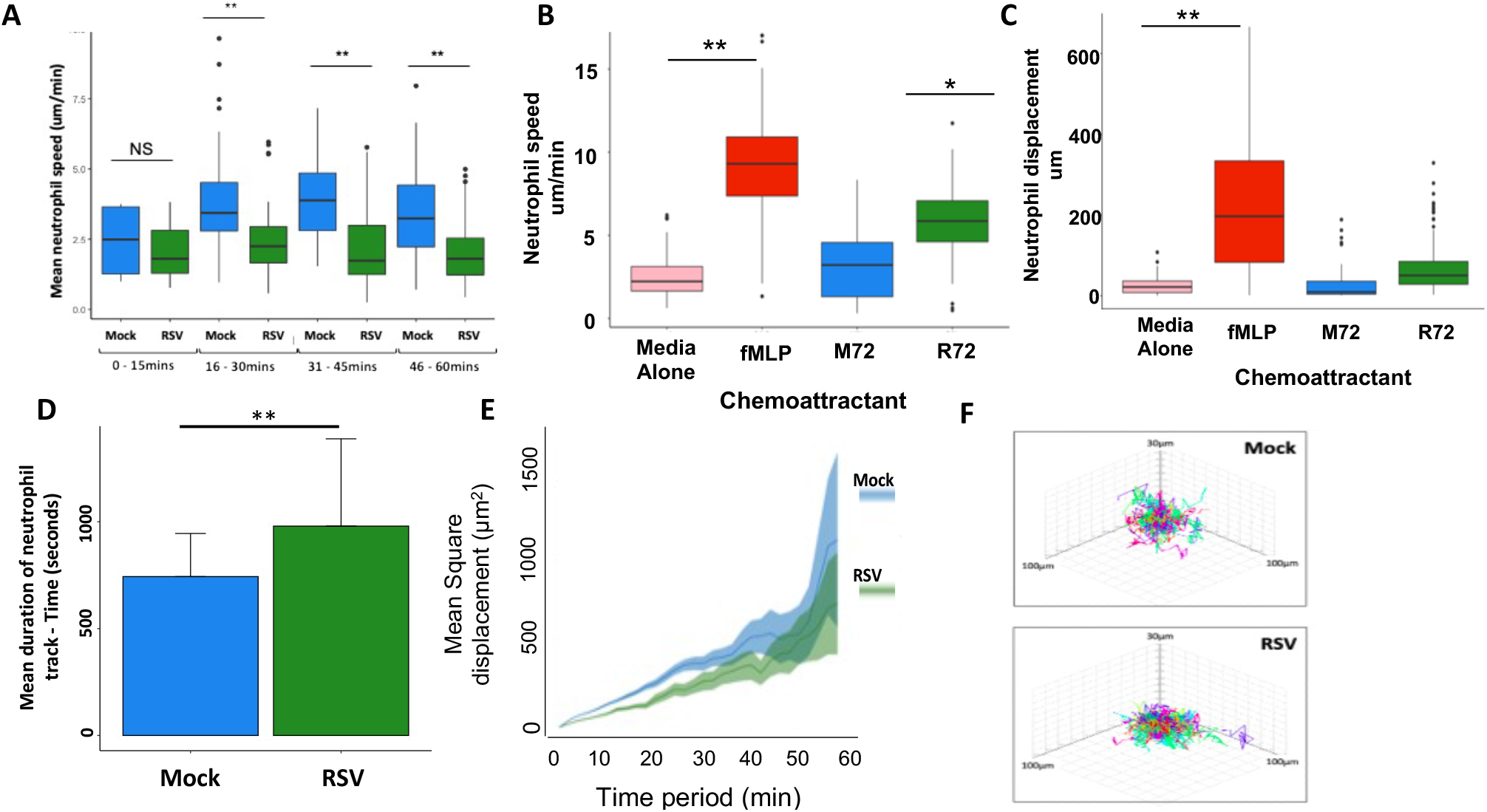
**A)** Average neutrophil speed during trans-epithelial migration across mock and RSV infected AECs. This time series was split into 15-minute segments. **B)** 2D Chemotaxis in ibidi chemotaxis chambers showing the average speed neutrophils move in response to apical supernatants from AECs. **C)** Neutrophil displacement – the gross distance travelled by individual neutrophils in response to apical supernatants from AECs. **D)** Average track duration of 200 individual neutrophils during movement through mock and RSV infected AECs. Speed of tracks were calculated using motilityLab. **E)** Mean squared displacement of neutrophils at each time point during movement across mock and RSV infected AECS. Statistical analysis between groups were performed using a Wilcoxon-Mann-Whitney U test, where significance was found this is indicated on the chart. *** p < 0.001. **F)** Visualisation of 50 tracks

To determine whether this was associated with chemoattractants in the apical supernatant, we tracked the movement of naive neutrophils towards apical supernatants collected from RSV or mock infected AEC cultures using specialised 2D chemotaxis chambers (**Figure 4B**). Here, we found that neutrophils exposed to supernatants collected from RSV infected AEC cultures moved faster (0.205 μm/sec ± 0.001) than neutrophils exposed to supernatant collected mock treated AECs (0.110μm/sec ± 0.001) *(p* < 0.05) (**Figure 4B**). At their fastest recorded speed (0.568μm/sec ± 0.049), neutrophils exposed to supernatants collected from RSV infected AECs were 1.5 times faster (*p* = 0.018) than those migrating toward supernatant collected from mock treated AECs (0.377μm/sec ± 0.014). There was no significant difference between the neutrophils exposed to media only.

#### DURATION

Measuring cumulative displacement (i.e., total distance travelled), we found that neutrophils exposed to supernatants collected from RSV infected AECs moved significantly (*p*= 0.016) further (349μm ± 49.9) than those exposed to media alone (182.7μm ± 8.96) (**Figure 4C**). We also measured the duration of neutrophil tracks during trans-epithelial migration (i.e., the length of time each neutrophil interacts with epithelium). Here we found that neutrophils interacted with RSV infected AECs (368μm ± 49.9) for longer than mock infected AECS (**Figure 4D**). We found no significant difference in linearity, or ‘track straightness’ between neutrophils observed migrating through RSV infected AECs or the mock (**Supplementary Figure).**However, neutrophils moving through mock infected AECs showed significantly (*p*<0.001) greater total displacement (137.19μm ± 4.51) compared to those moving through RSV infected AECs (107.05μm ± 4.86) (**Figure 4E**).

#### DIRECTIONALITY

We performed some preliminary analysis of the Z-tracks of neutrophils (*n*=20) migrating across RSV infected AECs (3 representative tracks **Supplementary Figure S4**). This showed preliminary evidence that neutrophils may move bidirectionally across the AECs, migrating to the apical side of RSV infected airway epithelium and returning to the basolateral compartment, as soon as 15 minutes after migration. This supports the hypothesis that the increased CD11b expression in neutrophils in the basolateral compartment is due to a proportion of those neutrophils having undergone reverse migration.

### Neutrophil clustering occurs 20 minutes after neutrophil recruitment

We previously observed that neutrophils adherent to RSV infected epithelium formed large clusters (22).Here, we aimed to quantify this clustering and determine whether it was significantly enhanced during RSV infection. To do this we first determined the expected nearest neighbour median distance, assuming an even distribution of the average number of adherent neutrophils. As more neutrophils are adherent to RSV infected AECs compared to mock infected, we determined that the median distance of the *expected* nearest neighbour was shorter in RSV infected AEC cultures (140μm ± 24.3) in comparison to neutrophils adherent to the mock infected epithelium (169.4μm ± 57.8). The *observed* nearest neighbour median distance was also shorter in RSV infected AEC cultures (61.32μm ± 30.9) in comparison to neutrophils adherent to the mock infected epithelium (119.4μm ±51.6) (**Figure 5B**). Importantly, the observed distances between neutrophils adherent to RSV infected AECs (61.32μm ± 30.9) was significantly (p<0.001) shorter than the expected distance (140μm ± 24.3) (**Figure 5B**). This indicates that the distribution of these adherent neutrophils on the epithelial surface is neither random nor uniform.

**Figure 5.**
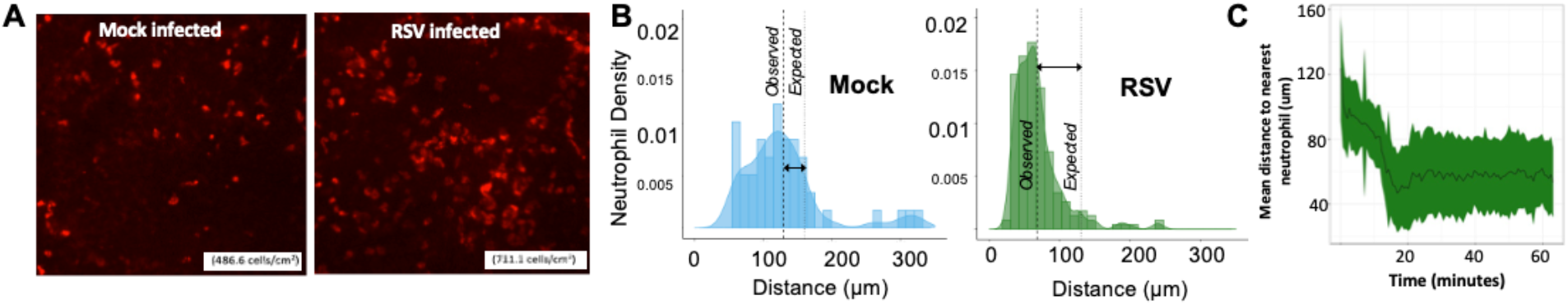
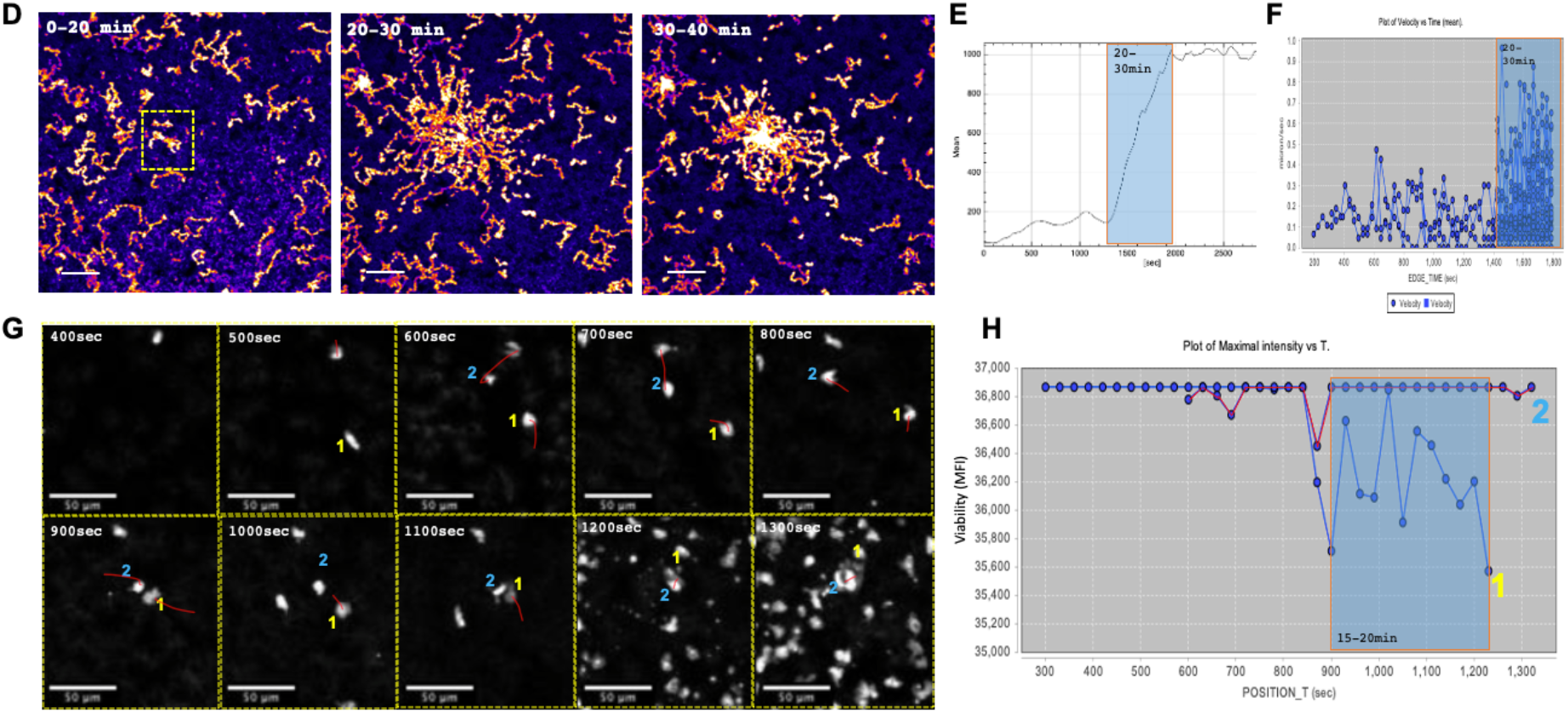
Neutrophil swarming dynamics during RSV infection in a human model. **A** Distribution analysis of neutrophils adherent to mock (left panel) and RSV (right panel) infected AECs following trans-epithelial migration. AECs were fixed after 1 hour; neutrophil position was processed using ImageJ and coordinates analysed using R version 4.0.3. **B)** Histogram showing the frequency of neutrophils the distribution of distance to nearest neighbouring neutrophil calculated from neutrophils adherent to the mock (blue) or RSV (green) infected AECs after 1 hour. Thin dashed lines show median distance to nearest neighbouring neutrophil. Thick dashed lines show median distance to nearest neighbouring neutrophil assuming a uniform distribution of adherent neutrophil counts. Median distances were compared using a Wilcoxon-Mann-Whitney U test. **C)** Average nearest neighbour distance of neutrophils over time during migration through RSV infected AECs, calculated as above from fast time-lapse video microscopy (n=200). Mean shown with dark green line. Range (min-max) indicated in colour block. Track metrics calculated using motilityLab. **(D)** Time-course of migration tracks on apical side of AEC following addition of neutrophils on basolateral side. Maximal intensity projection of the first 20- (left panel) or 10-minute (middle and right panels) time intervals. Scale bars, 100 μm. (**E**) Time-course of neutrophil accumulation (by MFI) on the apical surface of RSV infected AECs. (**F**) Time course of neutrophil accumulation by number of neutrophils on apical side per time-interval is shown. (**G)** images and MFI (**H)** of a single neutrophil (#1) becoming calcein red orange-negative at 15 min and its correlation to neutrophil-amplified chemotaxis (red tracks) Scale bars, 50 μm. Neutrophil positions were processed using ImageJ (imagej.nih.gov/ij/) and coordinates analysed using R version 4.0.3. (26)

We then determined when, during trans-epithelial migration, neutrophils begin to cluster to RSV infected AECs. This was defined as the earliest timepoint that the observed and expected nearest neighbour median distance became significantly different (**Figure 5C**). We found that after 20 minutes, the observed median (± interquartile range (IQR)) difference between neutrophils and their nearest neighbours (102.9μm ± 55.8) was significantly shorter (*p* < 0.001) than the distance expected by chance alone (140μm ± 24.3) (**Figure 5C)**. This suggests that neutrophil clustering begins to occur around 10 mins after the addition of neutrophils to the basolateral side of AECs in our model. Between 10-20 minutes we detected a more marked rate of decline in nearest neighbour distance to 42.9μm ±55.8. After this time phase, and between 20-60 minutes, the mean distance to nearest neutrophil stays constant around 50μm (**Figure 5C**).

### Location of neutrophil clustering is initiated by a dying neutrophil

In order to observe the initiation of neutrophil clustering to RSV infected AECs we used a higher-speed and resolution image capture system (see **Video 3**). Here neutrophils, labelled with a fluorescent viability stain (calcein red-orange), were found to rapidly nucleate and form a cluster on the apical side of the AECs approx. 20 minutes after the basolateral addition of neutrophils (**Figure 5DEF**). This pattern of clustering bears resemblance to previous *in vivo* work which described this observation as neutrophil swarming (24). Previous in vitro assays with bacteria have indicated that dying neutrophils precede the swarm-like formation of neutrophil clusters(25). To determine probable source of the nucleation point in our model, we segmented a region of interest immediately adjacent the cluster nucleus (shown in **Figure 5D**) and tracked the neutrophils in the preceding 0-20 minutes. This data (**Figure 5GH**) shows that the timing of the death (loss of viability stain) of a single neutrophil (labelled #1 in Figure 6GH) correlated to the amplified recruitment of the neutrophil population at 20 min, that was directed towards the unviable neutrophil (see **Video 4**). This indicates that neutrophil death may serve as a catalyst for swarming during RSV infection.

## Discussion

This study has, for the first time, performed spatial, temporal and functional analysis on migrating neutrophils in response to RSV infection of primary human airway epithelial cells. Our findings identify three putative phases of neutrophil migration:

1. Initial chemotaxis and adherence
2. Activation and reverse migration
3. Amplified chemotaxis and clustering

Firstly, we found that RSV infection led to greater, but slower, net movement of neutrophils across RSV infected AECs in comparison to mock infected AECs. This contradicted with our findings that, in the absence of AECs, neutrophils moved faster towards RSV infected supernatants (that contained elevated levels of the potent neutrophil chemoattractants CXCL8 (IL-8) and CXCL10 (IP-10) (**Supplementary Figure S3**), compared to neutrophils exposed to mock infected AEC supernatants. This suggests that an airway epithelial cell factor is responsible for slowing down the migration of neutrophils in the RSV infected AEC model. This could be due to the interaction of epithelial ICAM-1 to neutrophil integrin LFA-1, as we have previously shown (27). Alternatively, cell syncytia formation, a known histopathological characteristic of RSV infection, could increase epithelial impedance by reducing the availability of accessible cell-cell junctions through which migration may occur(27,28). This may not only slow down the initial neutrophil response to RSV infection, but also prolong its duration and slow its resolution, contributing to a heightened period of inflammation.

Secondly, we found that trans-epithelial migration led to apical (but not basolateral) secretion of soluble neutrophil granular factors including MMP9, myeloperoxidase (MPO), and neutrophil elastase (NE) that correlated to epithelial damage. Migration also led to greater expression of neutrophil activation markers CD11b, CD64, CD62L, NE and MPO, with highest expression recorded on neutrophils recovered from the apical compartment of RSV infected AECs. Interestingly, neutrophils recovered from *basolateral* compartments of the AEC model also increased expression of CD11b, CD64, CD62L and MPO, but not NE, compared to the no-AEC control. This is supported by clinical studies of children hospitalised with RSV bronchiolitis, which showed that neutrophils with upregulated CD11b were recoverable both from the airways by bronchiolar lavage and from peripheral circulation(9). Furthermore, neutrophils recovered from peripheral circulation of infants with RSV bronchiolitis have been shown to contain RSV mRNA(11). Viraemia due to RSV has not, to our knowledge, been observed clinically, which suggests that the virus causes symptoms because of its effects on the respiratory tract. This poses an important question: are activated neutrophils preferentially selected for migration or does the process of trans-epithelial migration increase expression of neutrophils activation markers? To address this, we cultured our AECs on membrane inserts with a smaller pore size (0.4μm), that prevented neutrophil migration. Here, following RSV infection, we did not find any difference in CD11b, CD64, CD62L and MPO expression on basolateral neutrophils compared to neutrophils exposed to media alone (no AECs). This suggests that neutrophils increase their expression of CD11b, CD64, CD62L and MPO during or following migration, rather than an existing subpopulation of activated neutrophils being selected for migration. This was different for NE. Here we detected a decrease in NE expression when neutrophils were exposed to basolateral side of RSV infected AECs, compared to mock infected. as a similar pattern in NE expression is seen when comparing neutrophils exposed to the no-AEC mock and RSV supernatants, suggesting that secreted factors may drive this change. Therefore, the presence of these ‘activated’ neutrophils on the *basolateral* side of the AECs following migration, may suggest that neutrophils, which are ‘activated’ by the apical environment, reverse migrate to the basolateral side of AECs (see Graphical Abstract). To investigate this further, we analysed the Z-tracks of neutrophils migrating across RSV infected AECs (**Supplementary Figure S4**). This showed that neutrophils can move bidirectionally across the AECs, quickly migrating to the apical side of RSV infected airway epithelium and returning to the basolateral compartment, as soon as 15 minutes after migration.

Next, we found that RSV infection led to greater numbers of neutrophils remaining adherent to RSV infected AECs. This may be due to increased expression of host cell receptors including ICAM-1 (intercellular adhesion molecule 1), which we (and others) have previously shown to be upregulated *in vitro* airway models of RSV infection, and mediate neutrophil adherence to RSV infected AECs (22) (27). In support of this, we found that fewer neutrophils remained adherent to mock infected AECs exposed to RSV infected supernatant placed apically (in comparison to RSV infected AECs). This suggests that properties of the epithelial cells, rather than the apical milieu of the RSV infected epithelium *per se,* are responsible for neutrophil adherence. Interestingly, neutrophils were observed to adhere in clusters to the apical surface of RSV infected AECs. Neutrophil clustering has been shown as a vital mechanism for host defence against pathogens(22,29). However, large clusters have been associated with inflammatory disease in both human and murine models(30,31). In models of RSV infection, increased neutrophil adherence and clustering has been associated with greater AEC damage (21,22,32,33).

To examine the formation of these clusters in more detail, we used higher-resolution timelapse microscopy to image the early movement of neutrophils moving across RSV infected AECs. This showed that clusters can result from the coordinated convergence of neutrophils, which resembled reports of neutrophil swarming in response to infection and sterile injury in *in vivo* models (23,24),(22). This pattern of neutrophil movement is thought to be initiated by the release of danger-associated molecular patterns (DAMPs) from neutrophils, which induce a transcriptional switch in their neighbours and coordinate a collective movement (34). LTB4 is a leukotriene DAMP, released by dying neutrophils and has been shown to mediate this directed leukocyte movement previously (24,30,35,36). We also showed that neutrophil death, or loss of cell viability, may serve as a catalyst for swarming during RSV infection. Here, we speculate that the formation of clusters of neutrophils, seen in our model, is not mediated by the RSV infected AECs, but instead due to neutrophils themselves triggering and maintaining clustering behaviour. The ability of neutrophils to affect others around them is an emerging area of research, and neutrophil quorum signalling has been shown to coordinate neutrophil collective responses to wound healing(23,24,37). Whether this response is a natural protective mechanism in countering an RSV infection or an aberrant response remains to be fully explained, but focal damage to AECs *in vivo* may allow increased neutrophil migration.

In summary, neutrophil activation has been shown as a key precursor to the development of severe respiratory symptoms in children with RSV bronchiolitis (38). Here we have shown that contact of neutrophils with AECs and/or trans-epithelial migration through RSV infected AECs is essential for upregulation of neutrophil activation markers, including CD11b expression. We describe distinct, measurable patterns of neutrophil movement, including the formation of neutrophil clusters on RSV infected AECs. We present evidence of the bidirectional movement of neutrophils across AECs during RSV infection and return of activated neutrophils to the basolateral side of infected AECs. This could explain how neutrophils with upregulated CD11b and presence of viral products were recoverable in blood neutrophils of babies with RSV bronchiolitis (39) (9) (11),(23,34,37) and could have important systemic implications for severe disease sequalae. This is a critical area for discovery and the model that we have developed here could be used to unravel important disease mechanisms, including the key question of how RSV accesses extra-pulmonary sites during infection. Future work should include the identification of new biomarkers of specific neutrophils sub-populations responsible for driving severe disease outcomes; screening anti-inflammatory drugs; and determining new mechanisms that may aid disease resolution.

## Materials and Methods

### Participants

Peripheral blood and airway epithelial cells were obtained from healthy adult donors at UCL GOS Institute of Child Health. Written informed consent was obtained from all donors prior to their enrolment in the study. Study approval was obtained from the UCL Research Ethics Committee (4735/002). All methods were performed in accordance with the relevant guidelines and regulations.

### Neutrophil Isolation

Venous blood was collected in EDTA (Ethylene Diamine Tetra Acetic) tubes (Greiner). Neutrophils were then ultra-purified using an EasySep Direct Neutrophil isolation kit (Stem Cell Technologies) according to the manufacturer’s instructions and subsequently stained with Cell Trace Calcein Red-Orange cell stain (ThermoFisher) and processed as described previously(40).

### Transepithelial Migration Model

This study modified the transepithelial migration model described by Herbert et al (2020) to utilise undifferentiated human AECs, grown at air liquid interface for 7 days as opposed to 28-day ciliated cultures. This ensures a flatter more uniform culture which is possible to live image using confocal microscopy(22). AECs cells were cultured on porous PET inserts (Greiner) with pore size of 3μm to allow migration or 0.4μm to prohibit it. AEC cultures were infected with RSV 24 or 72 hours prior to the addition of neutrophils. Mock infected AECs with RSV infected AEC supernatant were used as a control (RSV Sup), as were mock infected AECs with n-Formylmethionine-leucyl-phenylalanine (fMLP) 100nM placed apically as a positive control for neutrophil chemotaxis. 400μl of supernatant was added underneath the membrane insert for each experimental group, Mock, RSV, RSV supernatants only and fMLP control. Excess supernatant prior to neutrophil migration was stored at −20 for use in chemotaxis and activation experiments. Neutrophils were then added to the basolateral side of all membrane inserts, then left to incubate for 1 or 4 hours. After migration, neutrophils were collected from the apical side of the epithelial cells for quantification. Supernatants were collected and membrane inserts were fixed and stained. Mock infection was performed by inoculating with sterile media in the place of viral inoculum.

### Virus purification and quantification

Recombinant GFP tagged RSV A2 strain was kindly provided by Jean-Francois Eleouet and described in Fix et al(41). Viral stock preparation and quantification of viral titre was performed using HEp-2 cells (ATCC CCL-23) was performed as described previously(22).

### Microscopy

AEC cultures were fixed for microscopy using 4% paraformaldehyde (v/v) and mounted using 0.1M n-Propyll Gallate in glycerol:PBS (9:1). Images of fixed cultures were acquired using an inverted Zeiss LSM 710 confocal microscope using a 20x Plan Achromat LWD objective. Live cell imaging was performed using a Perkin Elmer UltraView spinning disk (CSU22) confocal microscope. All microscopes were located in the UCL GOS Institute of Child Health Imaging Facility (London, UK).

### Quantification of migrated and adherent neutrophils

The number of migrated neutrophils was quantified as described previously (22). Flow cytometric analysis of CD11b, NE, MPO, CD64 and CD62L expression was performed as described previously(21). Antibodies used are provided in supplementary information. Image acquisition was the same as above. Neutrophils were counted using an ImageJ counting tool.

### Quantifying neutrophil distribution

To investigate whether neutrophil adherence to AECs was uniform (null hypothesis =0) or whether they clustered, a method of measuring neutrophil distribution was required. Images taken of AECs after 1 hour of migration were taken as described previously, then analysed using ImageJ to first identify neutrophils within the image, determine their 3D coordinates. These coordinates were then exported into R and analysed using the R package spatstat to then calculate the distance between the centre-point of each neutrophil and the centrepoint of its five nearest neighbours (42). However, it is expected that if more neutrophils are adherent to RSV infected AECs in comparison to mock, that they would be closer together by crowding alone. To account for this, the distance apart each neutrophil would be if neutrophils were evenly spread over the AEC area was calculated from the numbers of neutrophils counted as adherent to the same AECs. ImageJ macro code for determining 3D coordinates and subsequent R code for distance analysis is available upon request.

### Analysis of neutrophil chemotaxis

Time-lapse videos were analysed using Icy (https://icy.bioimageanalysis.org/), programmed to run a spot detection protocol for each frame of the stack, detecting light objects close to 10μm diameter on a dark background to identify neutrophils within the image. Then, a spot tracking plugin was used to map the displacement of each neutrophil through each time point image (43). The tracks produced were then visually checked for tracking accuracy and where erroneous tracks were found, corrected manually, or removed from analysis. Raw spot files and track files were exported for further analysis using R. Data sets detailing total displacement, net displacement, average, maximum and minimum speed, and linearity were exported combined with replicate data sets for analysis using R and GraphPad Prism v8.0.

### Time-lapse imaging of neutrophil trans-epithelial migration in 4D (XYZT)

A black plastic 24 well plate with a glass coverslip bottom (Greiner) was used in place of a low binding 24 well plate (Greiner) to allow for live imaging of AEC cultures during transepithelial migration. Neutrophils were identified using Icy and a spot detection protocol initiated for each frame of the stack, detecting light objects close to 10μm diameter on a dark background. Then each spot was tracked using the spot tracking plugin using a 4D tracking algorithm (43). Tracks were then visually checked for tracking accuracy and where erroneous tracks were found, were either corrected manually or removed from analysis. Raw track coordinates were exported for further analysis using R package Motility Lab (44). Data detailing displacement, speed and linearity were exported. Analysis of neutrophil movement and distribution was performed using the R packages ‘motility lab’, ‘spatstat’ and ‘dyplr’ (45–47).

## Acknowledgements

LR was recipient of a Newton fellowship from The Academy of Medical Science (ref NIF004/1012). RLS was supported by the Great Ormond Street Children’s Charity (grant code W1802). CMS is a recipient of grants from Animal Free Research UK (AFR19-20274), BBSRC (BB/V006738/1), GOSH Children’s charity (COVID_CSmith_017) and the Wellcome Trust (212516/Z/18/Z). This research was supported by the NIHR Great Ormond Street Hospital Biomedical Research Centre. Microscopy was performed at the Light Microscopy Core Facility, UCL GOS Institute of Child Health supported by the NIHR GOSH BRC award 17DD08.The views expressed are those of the author(s) and not necessarily those of the NHS, the NIHR or the Department of Health. We thank Dr Shyam Sawhney for technical support and the healthy volunteers who donated airway cells and blood samples for this study

## Author contributions

All authors declare no conflicts of interest.

ER conceived and designed the study, conducted experiments, analysed data, and prepared the manuscript. JAH designed the study, conducted experiments, analysed data, and reviewed the manuscript. MP assisted with flow cytometry data analysis and review of the manuscript. LR, IL, and AP assisted with data collection and review of the manuscript. DM assisted with microscopy acquisition and reviewed the manuscript. MC-B conducted statistical analysis and reviewed the manuscript. RLS and CMS oversaw the funding application and contributed to study conception and design, data analysis and interpretation, and the write-up of the manuscript.

