## Supplementary Information for "Trans-epithelial migration is essential for neutrophil activation during RSV infection"

**FIGURE S1 (A**) Plotted XYZ coordinates of tracks of neutrophils moving through Mock (left) and RSV (right) infected airway epithelial cells (AECs) with start point centred on the origin. (**B**) Plotted XY coordinates of tracks of neutrophils moving towards airway supernatant collected from either mock (left) or RSV (right) infected AECs in a 2D chamber slide system with start point centred on the origin. (**C**) Speed of naive neutrophils exposed to apical supernatants from mock (blue) and RSV infected (green) AECs in a 2D chamber slide system in absence of AECs, compared to media alone and positive control fMLP. Image taken every 30 seconds for 1 hr and speed calculated using icy. (**D**) – Displacement of naive neutrophils exposed to supernatants from infected AECs in a 2d system in absence of AECs. Images taken every 30seconds for 1 hr. Statistical analysis between groups were performed using a Mann Whitney U test with Wilcoxon rank, where significance was found this is indicated on the chart. * p<0.05.

**A**

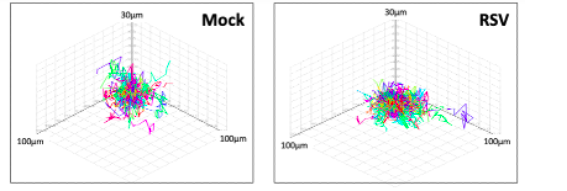

**B**

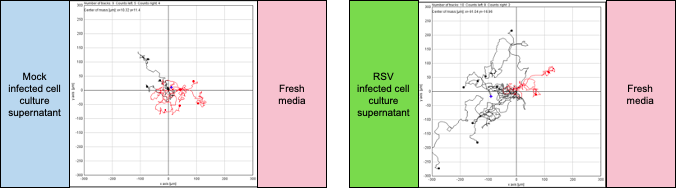

**C**	**D**

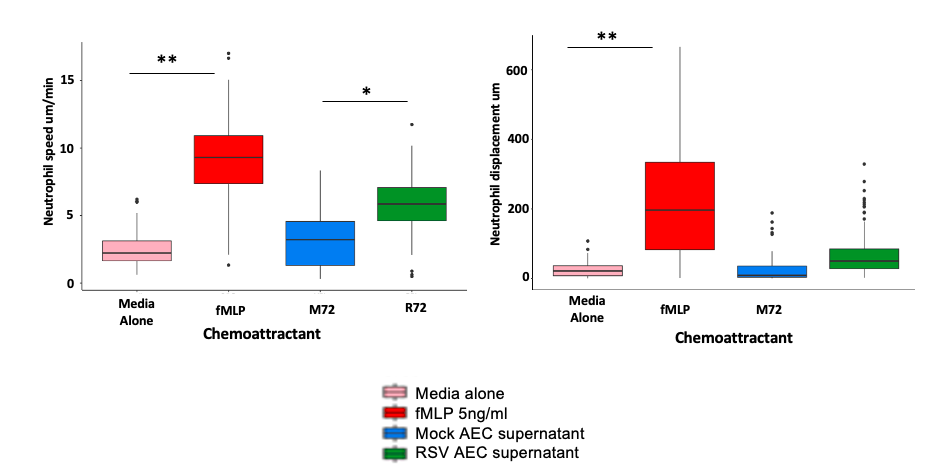

**Figure S2 –AECs grown on membrane inserts with a 0.4um pore size do not permit neutrophil interaction with AECs.** Z-projection of a single channel (calcein-red orange) z-stack image captured after 1-hour transepithelial migration experiment using AECs grown on 0.4um pore Thincerts (Griener). The level of the PET membrane of the insert is indicated with the yellow dashed line and neutrophils (white) are visible above but not below. Images taken using a confocal Zeiss spinning disk CSU22 system.

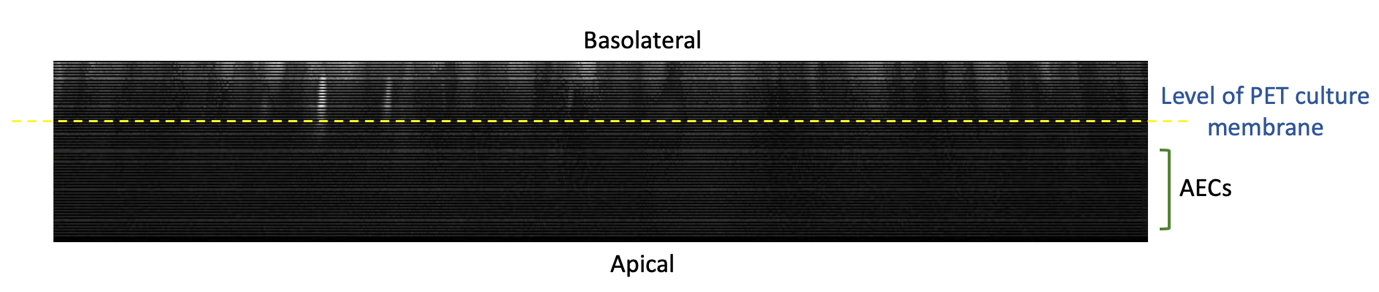

**Figure S3 - Concentration of IL-8 and IP-10 in the apical supernatants collected from RSV infected AECs**. Concentration of CXCL8 (IL-8) (left) and CXCL10 (IP-10) (right) in apical supernatants collected 1 hour's incubation of fresh media with AECs infected with RSV (green) for 24 or 72 hours or Mock (blue) infected AECs infected for 24 or 72 hours. IL-8 and IP-10 concentration were determined using a commercial ELISA kit (ThermoFisher). Groups were compared with One-Way ANOVA with Tukey adjustment for multiple testing. *= p<0.05. **=p<0.002.

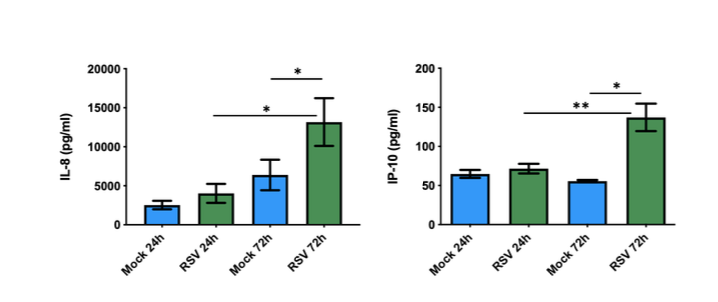

**Figure S4 - Track trajectories in the z-axis of 3 independent neutrophils migrating through RSV infected AECs.** Vector coordinates showing only the z position (x-axis) over time (y-axis) of three selected neutrophil tracks from a dataset of 200+ neutrophils migrating across AECs infected with RSV for 72hrs. Dotted line represents the position of the PET membrane, the top representing the basolateral and bottom the apical compartments.

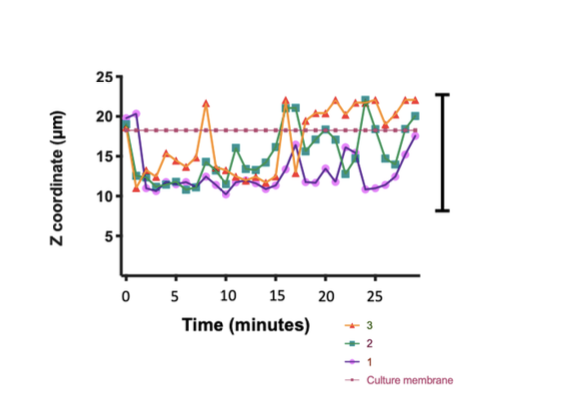
